# An orthogonal TRAP enables intersectional genetic access to activated neurons in the mouse brain

**DOI:** 10.64898/2025.11.30.691330

**Authors:** Nikolaos Chatziris, Brooke C. Jarvie, Can Liu, Zachary A. Knight

## Abstract

The study of neural circuits has been greatly enabled by methods for obtaining genetic access to activated neurons. However, these approaches typically tag neurons based on their response to only a single stimulus, which limits the ability to define precise subpopulations of cells. Here we describe an approach (X-TRAP) in which the activity-dependent expression of Flp recombinase is gated by branaplam, a small molecule that triggers splicing of the X-ON switch. We show that X-TRAP knock-in mice exhibit undetectable Flp recombination in the absence of drug and that branaplam treatment results in robust induction of recombination selectively in neurons that express FOS. Moreover, we show that X-TRAP is orthogonal to the widely-used TRAP system, such that these two approaches can be used in the same animal to label cells with Cre and Flp recombination in response to two different stimuli. We apply this strategy to map neural circuits that control food intake. This approach for intersectional, activity-dependent genetic labeling should enhance our ability to identify the neural correlates of behavior.

## Introduction

A basic goal of neuroscience is to link the activity of specific neural cell types to the functions of the brain. This effort has been greatly enabled by the ability to gain genetic access to functional populations of neurons so that they can be monitored and manipulated (Luo et al., 2018).

Marker genes provide one way to identify and access neural cell types (Gong et al., 2003; Lein et al., 2007). However, for many brain functions, there is no marker gene known that selectively labels the relevant cells. In these situations, an alternative approach is to genetically tag neurons based on their acute response to a stimulus (DeNardo and Luo, 2017; Franceschini et al., 2020). This can be achieved by coupling a biochemical correlate of neural activation, such as calcium influx or *Fos* expression, to the stable expression of a transgene, such Cre recombinase.

Several strategies for activity-dependent genetic labeling have been described (DeNardo et al., 2019; Guenthner et al., 2013; Kawashima et al., 2013; Lee et al., 2017; Sakurai et al., 2016; Sørensen et al., 2016; Wang et al., 2017). However, the most widely-used approach is Targeted Recombination in Active Populations (TRAP), which is based on a transgenic mouse in which a tamoxifen-regulated Cre recombinase (CreER^T2^) has been targeted to the *Fos* locus (Allen et al., 2017; DeNardo et al., 2019; Guenthner et al., 2013). As a result, CreER^T2^ is expressed in neurons that are acutely activated by a stimulus, and Cre recombination can be induced in these cells by peripheral injection of 4-hydroxytamoxifen (4-OHT). Relative to other approaches, TRAP has the advantage that it enables genetic labeling of activated neurons throughout the brain in a non-invasive and straightforward way (Franceschini et al., 2020).

Despite their utility, TRAP and related approaches have a limited ability to define precise subpopulations of activated cells. This is because, at any given time, *Fos* is expressed in multiple populations of neurons throughout the brain, and only a fraction of these *Fos+* cells are expressed as a response to the intended stimulus. Moreover, even among the stimulus-responsive cells, heterogeneity arises from the fact that there are multiple components to the brain’s response to any stimulus (e.g. sensory, motor, affective). Thus, there is a need for methods that enable more fine-grained genetic access to functional populations of neurons.

We reasoned that the specificity of approaches like TRAP might be improved if it were possible to label the neurons activated by two different stimuli administered on two different days. This would enable, for example, genetic access to only the cells that represent the intersection of two different activation patterns and therefore share a common feature. Here we describe a strategy for doing this in which the activity-dependent expression of Cre and Flp recombinases is gated by two different small molecules.

## Results

We sought to develop an approach in which the expression of Flp recombinase in the brain is (1) activity-dependent, (2) gated by the peripheral administration of a small molecule, and (3) regulated orthogonally to the TRAP system, so the two approaches can be used in the same animal. To do this, we took advantage of the fact that the small molecule branaplam (Cheung et al., 2018; Keller et al., 2022) can induce alternate splicing of a small cassette known as the X-ON switch (**Fig 1a**) (Monteys et al., 2021). We fused this X-ON cassette to the coding sequence for Flp recombinase and then generated knock-in mice in which this transgene was targeted to the *Fos* locus, so that X-ON-Flp expression is controlled by the endogenous *Fos* regulatory elements. This transgene was designed such that, in the absence of branaplam, the resulting mRNA lacks an in-frame start codon and therefore does not produce functional Flp protein (**Fig. 1a**). Administration of branaplam triggers alternative splicing that introduces an in-frame start codon and allows Flp expression (**Fig. 1a**). We refer to this approach in which Flp-dependent TRAP is controlled by the X-ON switch as “X-TRAP.” Importantly, because branaplam has no effect on CreER^T2^, and tamoxifen has no effect on the X-ON switch, TRAP and X-TRAP can be used together in the same animal to enable intersectional genetic access to the neurons that respond to two different stimuli (**Fig. 1b**).

**Figure 1.**
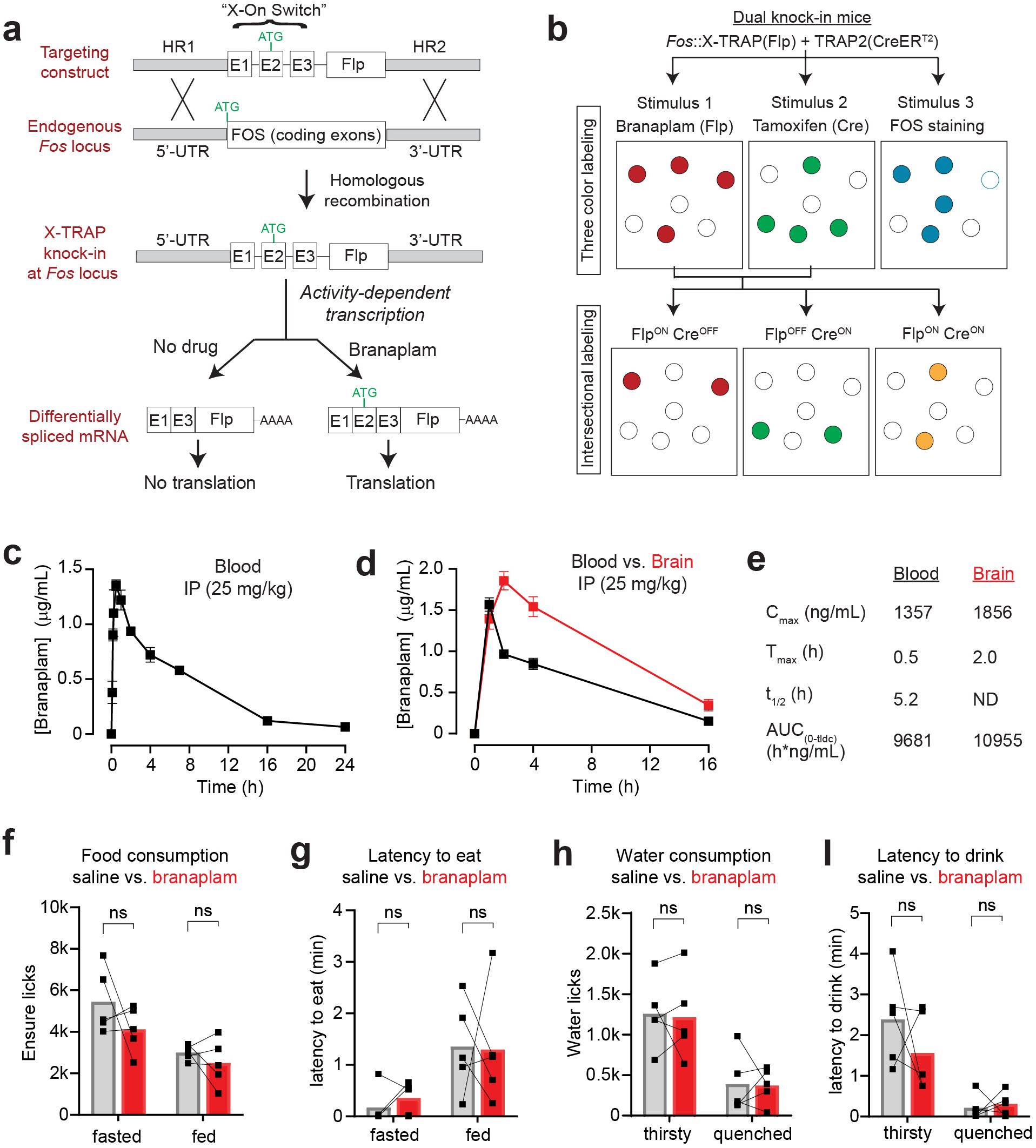
Activity-dependent Flp recombination using X-TRAP. **a.** Schematic describing design of the X-TRAP knockin mouse, in which a cassette containing the "X-On Switch" and FlpO is knocked in immediately upstream of the start codon of the *Fas* gene. Activity-dependent transcription at the *Fas* locus produces FlpO mRNA that is not translated into protein due to lack of an in-frame start codon. Treatment with branaplam causes alternative splicing to produce a transcript that contains a start codon and is translated. **b.** The X-TRAP system, when used in conjunction with the TRAP2 allele, enables intersectional labeling of activated neurons in response to sets of stimuli. Top: Neurons expressing FOS in response to three different stimuli are visualized by labeling neurons with X-TRAP, TRAP2, and immunostaining for FOS. Bottom: When used in conjunction with reporter mice or viruses, this enables genetic access to cells that are the intersection of neurons activated by different stimuli. **c.** Concentration of branaplam in the blood after IP injection (25 mg/kg). **d.** Concentration of branaplam in the blood and brain after IP injection (25 mg/kg). **e.** Pharmacokinetic parameters for branaplam in blood and brain. **f-1.** IP injection of branaplam (25 mg/kg) has no effect on food intake **(f),** latency to eat **(g),** water consumption **(h),** or latency to drink **(i).** Data are mean ± sem.

We first confirmed that branaplam is compatible with in vivo experiments in behaving animals. Intraperitoneal (IP) injection of branaplam (25 mg/kg) resulted in peak concentrations in the blood at 30 min (1357 ± 49 ng/mL, or 3.4 µM), which then gradually declined with a half-life of ∼5 hours (**Fig. 1c**). Peak concentrations in the brain (1856 ± 111 ng/mL, or 4.6 µM) were slightly delayed relative to the blood (T_max_ ∼2 hours) and then declined with a similar half-life (**Fig. 1d,e**). For comparison, 4-OHT has a blood half-life of ∼30 hours after IP injection (Zhong et al., 2015). Thus, these data suggest that branaplam could be used to tag neurons that are activated in vivo within a window of several hours.

Short-term oral administration of branaplam is well-tolerated (Monteys et al., 2021), but it is unknown whether IP injection of branaplam has toxicity or effects on behavior. We administered escalating doses of branaplam and did not observe any morbidity and effects on eating or drinking up to 25 mg/kg (**Fig. 1f-i**). In contrast, there were signs of sickness in some animals after injections at 50 mg/kg. We therefore used 25 mg/kg for the studies that follow.

### X-TRAP exhibits a low level of recombination at baseline

We crossed the *Fos^X-TRAP^* mouse to a reporter line that expresses nuclear-localized Tomato after Flp recombination (*Fos^XTRAP/+^ Igs7^CAG-FSF-Tomato, CAG-LSL-GFP^*) so that we could visualize X-TRAP activity throughout the brain (**Fig. 2a**). In the absence of branaplam, essentially no FLP recombination was detectable in the brains of adult mice (no Tomato+ cells observed in brain slices from three animals; **Fig. 2b** and **Fig. S1a**). This indicates that functional FLP expression is absolutely dependent on branaplam-induced splicing.

**Figure 2.**
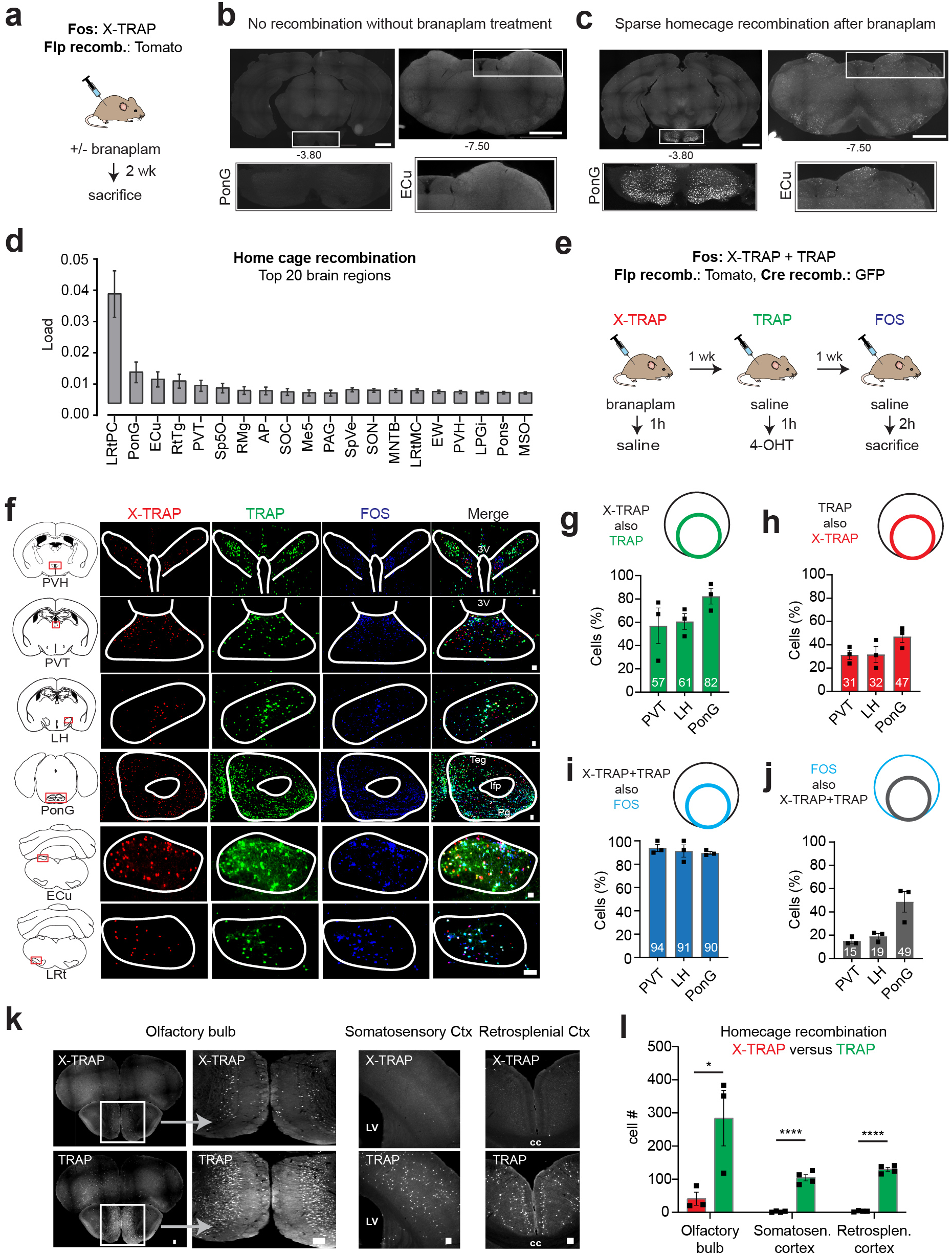
Baseline recombination with X-TRAP is low and similar to TRAP. **a.** Schematic for measuring homecage recombination with X-TRAP. **b.** There is no detectable Flp recombination in X-TRAP mice without branaplam treatment. Two coronal sections are shown with insets of pons (PonG) and external cuneate (ECu) nuclei. **c.** Homecage recombination after branaplam treatment in coronol sections matched to panel b. **d.** Top 20 regions with most homecage recombination in branaplam treated mice. **e.** Schematic for comparing baseline recombination between XTRAP and TRAP. **f.** X-TRAP and TRAP recombination, and endogenous FOS expression, in six brain regions. **g.** % X-TRAP+ cells that are also TRAP+. **h.** % TRAP+ cells that are also X-TRAP+ i. % dual labelled (X-TRAP+ and TRAP+) cells that are also FOS+. **j.** % FOS + that are also labelled by both X-TRAP and TRAP. **k.** Three brain regions that show higher baseline recombination with TRAP than X-TRAP. I. The number of cells labelled in the brain regions from k. Scale bars = 1 mm (panels band c) or 0.1 mm (panels f and k). NS, *P<0.05, **P<0.01, ***P<0.001, ****P<0.0001. Data are mean± sem.

We next measured the level of FLP recombination induced by branaplam injection at baseline (i.e. “home cage recombination”). Mice were given injection of branaplam (25 mg/kg) and then two weeks later tissue was processed for histology. We found that branaplam treatment induced sparse recombination in a subset of structures throughout the brain (**Fig. 2c,d** and **Fig. S1a**). To quantify this background recombination, we performed brain-wide imaging and used an automated pipeline for image registration and quantification to align this labeling to the reference mouse brain atlas (Yates et al., 2019). The 20 structures with the highest background labeling are show in **Fig. 2d**. Of note, many of these structures are involved in sensorimotor processing and likely to be activated during homecage activity.

To determine whether this background recombination represents specific labeling of neurons that are activated at baseline, we next directly compared the recombination induced by X-TRAP and TRAP, as well as expression of endogenous FOS, in the same animal. To do this, we generated by breeding mice that contain both the X-TRAP and TRAP2 alleles of *Fos* (TRAP2 is hereafter referred to as “TRAP”) as well as a reporter that expresses Tomato in response to Flp recombination and GFP in response to Cre recombination (*Fos^XTRAP/TRAP2^ Igs7^CAG-FSF-Tomato, CAG-LSL-GFP^*). These mice were subjected to a protocol (**Fig. 2e**) in which they received (1) injection of branaplam followed one hour later by injection of saline; (2) one week later, injection of saline followed one hour later by injection of 4-OHT; and (3) one week later, injection of saline followed two hours later by sacrifice, to enable subsequent staining for endogenous FOS. Saline injection was used to mimic a “vehicle injection” (the baseline state for many experiments), and the timing of 4-OHT and branaplam injection relative to saline was selected based on pilot experiments showing that this timing yielded the highest recombination for each method (see *Discussion*). Brain tissue was then stained and imaged for expression of Tomato (X-TRAP recombination), GFP (TRAP recombination) and endogenous FOS.

We found that X-TRAP and TRAP induced recombination in a sparse and mostly overlapping set of structures throughout the brain **(Fig. S1b**). Moreover, almost all of these structures also expressed endogenous FOS in a similar anatomic pattern (six examples are shown in **Fig. 2f**). To quantify the overlap between these markers, we measured the co-localization between X-TRAP, TRAP, and FOS in three brain regions that showed baseline labeling (PVT, LH, and PonG; **Fig. 2f-i**). This revealed that most of the neurons labeled by X-TRAP were also labeled by TRAP (57-82% of X-TRAP+ also TRAP+; **Fig. 2g**) and many TRAP+ were also X-TRAP+ (31-47%; **Fig. 2h**). Thus, these two methods capture an overlapping population of cells even when the labeling is performed on different days and in the absence of any specific stimulus.

Among the cells labeled by both X-TRAP and TRAP, a very high percentage also expressed endogenous FOS (91-94%, **Fig.2i**), confirming that these dual-labeled cells are indeed neurons activated at baseline. On the other hand, only a minority of all FOS+ cells were dual labeled by both X-TRAP and TRAP (15-49%; **Fig. 2j**). These observations are consistent with the notion that there is day-to-day fluctuation in the ensemble of cells that express FOS at baseline, and the dual-labeled cells (X-TRAP+ and TRAP+) are those that express FOS+ most consistently or at the highest level. Overall, these data show that (1) TRAP and X-TRAP label an overlapping population of neurons at baseline, and (2) most of these labeled neurons are activated, as measured by FOS expression.

While TRAP and X-TRAP induced similar baseline recombination in most brain regions, a few structures had much higher labeling by TRAP. These included the olfactory bulb and several cortical structure, such as somatosensory cortex and retrosplenial cortex (**Fig. 2k,l**). This difference may reflect the greater efficiency of Cre versus Flp recombinase, which allows TRAP to better detect low or transient FOS expression. Consistent with this, TRAP often captured a somewhat higher percentage of X-TRAP cells (**Fig. 2g**) than vice versa (**Fig. 2h**). This suggests there may be a trade-off between efficiency, which under some conditions is better with TRAP, and low background, which under some conditions is better with X-TRAP (see *Discussion*).

### X-TRAP enables Flp recombination in neurons activated by diverse stimuli

We next tested whether X-TRAP could be used to detect neurons activated by an acute stimulus. X-TRAP reporter mice were injected with branaplam and then, one hour later, challenged with one of three stimuli (**Fig. 3a**): (1) hypertonic saline, which activates neurons involved in fluid balance (Rinaman et al., 1997), (2) ghrelin, a hormone which activates neurons that promote hunger (Hewson and Dickson, 2000; Traebert et al., 2002; Wang et al., 2002), and (3) haloperidol, a dopamine D2 receptor antagonist which activates neurons in the midbrain dopamine pathway (Robertson and Fibiger, 1992). As a control, we also labeled the neurons activated by a vehicle injection (isotonic saline). Two weeks later, tissue was processed for histology and Tomato expression quantified across the brain.

**Figure 3:**
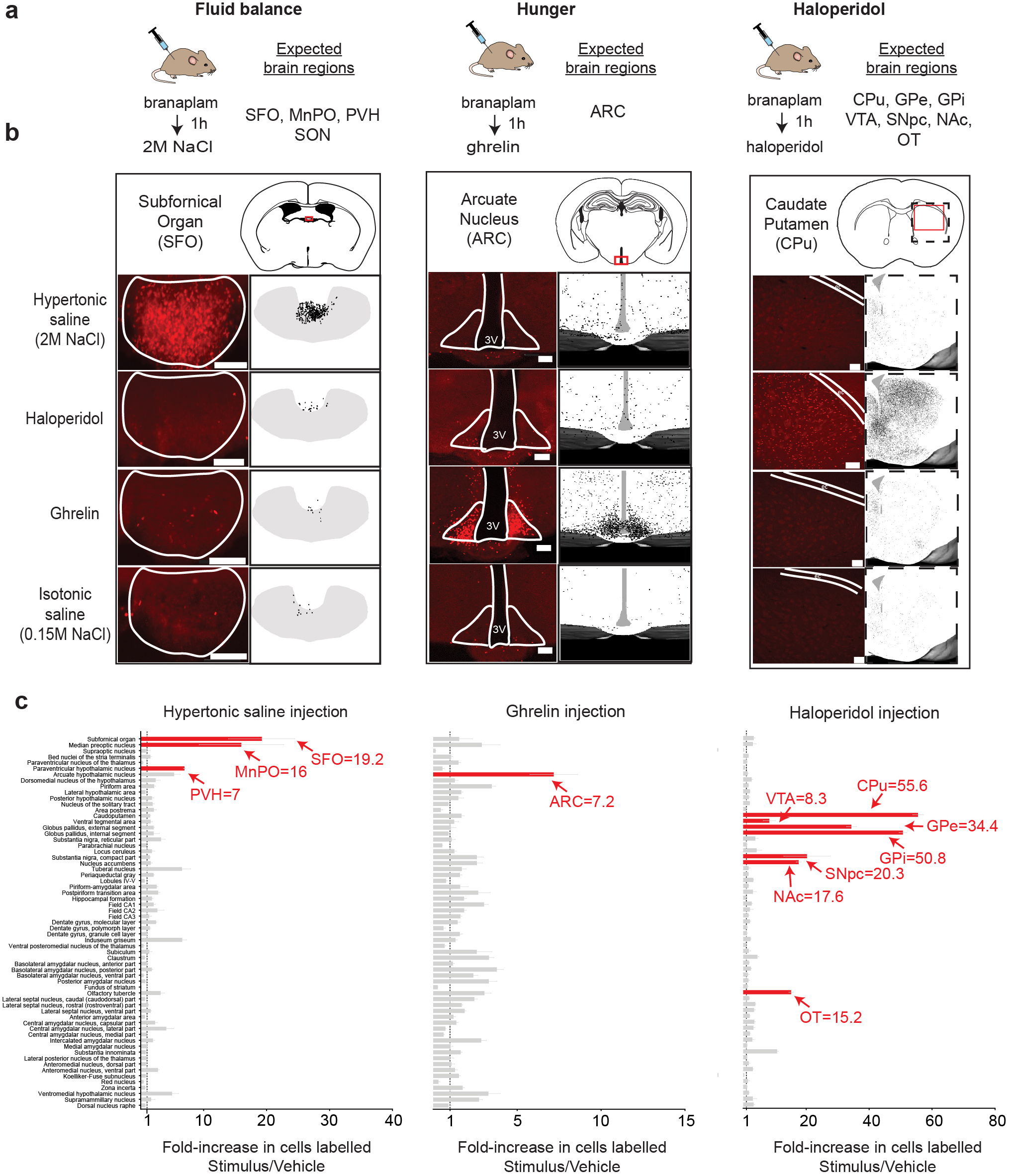
X-TRAP enables selective Flp recombination in neurons activated by diverse stimuli. **a.** Fos><TRAP was crossed to 1e ai224 reporter line, in which Flp recombination induces nuclear expression of Tomato. Mice were injected with branaplam (25 1g/kg, IP) and an hour later challenged with one of the following stimuli: Hypertonic saline (2M NaCl, 0.25 ml), ghrelin (2 1g/kg), haloperidol (-2 mg/kg), or control saline (0.15M NaCl, 0.1 ml). Mice were sacrificed two weeks later and Tomato xpression was quantified throughout the brain. **b.** Tomato expression of the SFO, ARC, and CPu in response to the indicated timuli. (Left) Example image from one mouse in red. (Right) Overlay of Tomato positive cells from n=4 mice. **c.** Tomato xpression in 63 midbrain and forebrain structures response to the stimuli indicated. Data are presented as the ratio of labeled ells compared to control (stimulus/vehicle). Key brain regions known to respond the indicated stimulus are highlighted in red.;cale bars are 0.1 mm. Data are mean± sem.

We found that X-TRAP reliably labeled the key brain regions known to be activated by each of these stimuli. For example, hypertonic saline injection caused strong labeling in the subfornical organ (SFO), a region critical for thirst (**Fig 3b**, left); ghrelin induced recombination in the arcuate nucleus (ARC), which contains ghrelin-sensitive neurons that control hunger (**Fig. 3b**, center); and haloperidol induced recombination in the caudate putamen (CPu), which is rich in the D2R-expressing neurons targeted by this drug (**Fig. 3b**, right). None of these brain regions showed strong labeling following vehicle injection, confirming that these responses are stimulus-specific.

To characterize the specificity of this labeling in an unbiased manner, we performed automated analysis of Tomato expression across 63 brain regions (Yates et al., 2019) and then quantified the Tomato expression in response to the stimulus relative to vehicle control (**Fig. 3c**). As expected, the principal brain regions known to be activated by each stimulus (**Fig. 3a**) also emerged as the most strongly labeled brain regions in this analysis (**Fig. 3c**). For example, the top three brain regions labeled by hypertonic saline were the SFO, median preoptic nucleus (MnPO) and paraventricular hypothalamus (PVH), all of which are critical for fluid balance (**Fig. 3c**, left); the top brain region labeled by ghrelin injection was the ARC (**Fig. 3c**, middle); and the top seven brain regions labeled by haloperidol were all structures in the dopamine pathway known to be activated by this drug (Robertson and Fibiger, 1992) (**Fig 3c**, right). The increase in cells labeled in these brain regions (stimulus relative to vehicle control) ranged from 7-fold (ghrelin/ARC) to 55-fold (haloperidol/CPu). The only brain region that we expected to detect as enriched, but did not, was the supraoptic nucleus (SON), which is activated by hypertonic saline (Carter and Murphy, 1990; Miyata et al., 1994). However, there was strong labeling of SON in the vehicle controls, likely due to stress caused by injection (Miyata et al., 1995), and this diminished the fold-enrichment. Overall, these data show that X-TRAP can be used to induce selective Flp recombination in neurons that respond to diverse stimuli throughout the brain.

### X-TRAP and TRAP label overlapping neurons in response to the same stimulus

A prerequisite for using X-TRAP and TRAP for intersectional labeling experiments is to establish that, in response to the same stimulus, these two approaches label the same neurons. To test this, we measured labeling in response to injection of the weight-loss drug semaglutide, which activates neurons that control food intake across multiple brain regions (Hansen et al., 2021). Mice containing both the TRAP2 and X-TRAP alleles as well as the Cre/Flp dual reporter (*Fos^XTRAP/TRAP2^ Igs7^CAG-FSF-Tomato, CAG-LSL-GFP^*) were subjected to a protocol in which they received three injections of semaglutide, or vehicle, each separated by a week, for labeling by X-TRAP, TRAP and endogenous FOS (**Fig. 4a**). We then characterized the expression of these markers in four brain regions known to express FOS following semaglutide treatment (Hansen et al., 2021): the area postrema (AP), nucleus of the solitary tract (NTS), parabrachial nucleus (PBN), and central amygdala (CeA). Of note, we performed this experiment in mice that were fasted (**Fig. 4**) as well as in mice that were ad libitum fed (**Fig. S2**) prior to semaglutide injection, and we obtained similar results.

**Figure 4.**
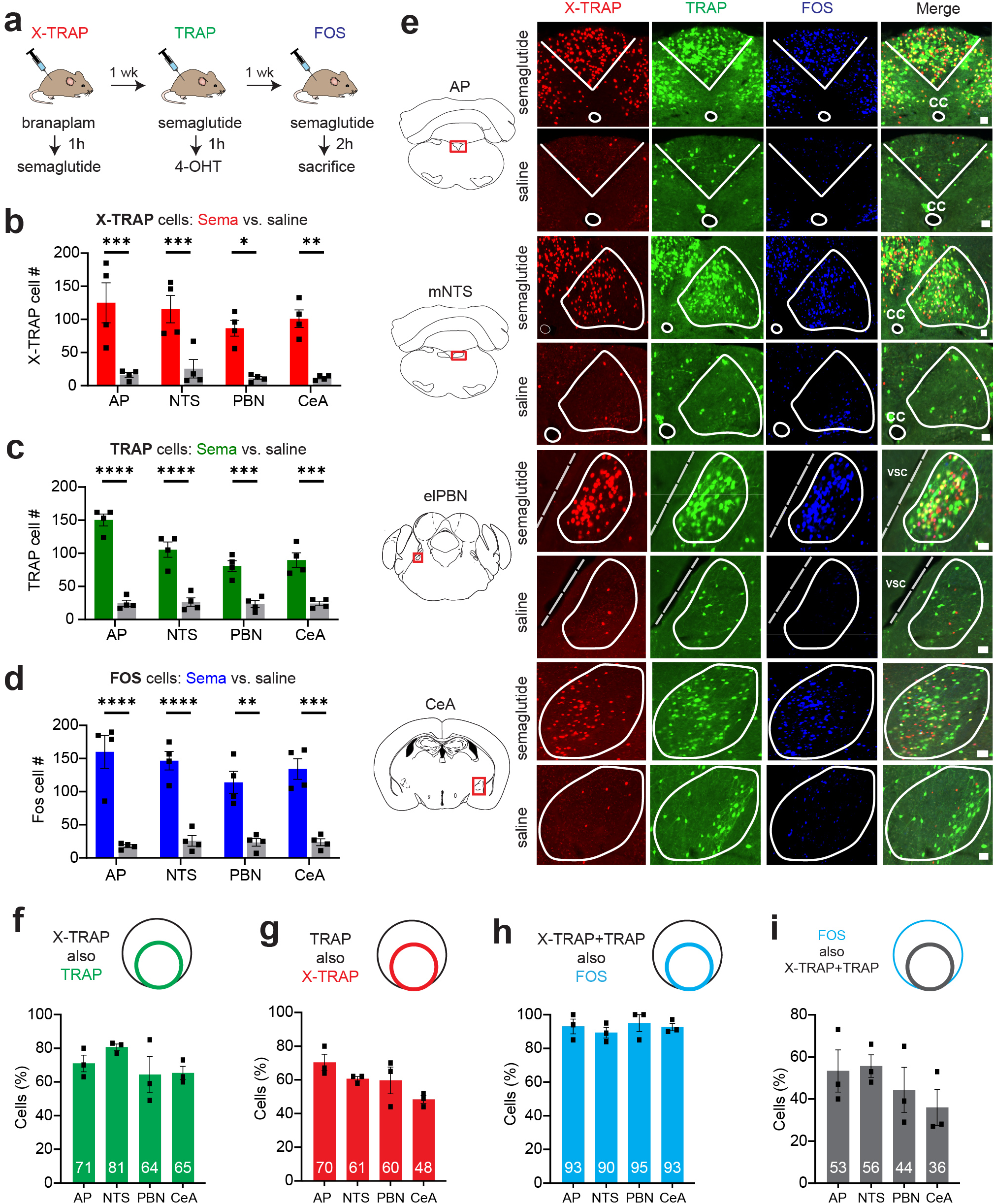
X-TRAP and TRAP label overlapping neurons in response to semaglutide. **a.** Protocol in which mice were challenged with semaglutide on different days, and activated neurons were labelled. **b-d.** Number of cells labelled in response to semaglutide or vehicle injection using X-TRAP **(b),** TRAP (c) or FOS staining **(d). e.** Examples of labelling in response to semaglutide and vehicle across the four brain regions shown. **f-i.** Overlap between markers following semaglutide injection. Panels show the percentage of cells labelled by X-TRAP also labelled by TRAP **(f),** TRAP also labelled by X-TRAP **(g),** X-TRAP+TRAP also labelled by FOS **(h),** FOS also labelled by X-TRAP+TRAP **(i).** Scale bars are 0.1 mm. NS, *P<0.05, **P<0.01, ***P<0.001, ****P<0.0001. Data are mean ± sem.

We found that semaglutide treatment caused a significant increase in the number of cells labeled by X-TRAP (**Fig. 4b,e**) and TRAP (**Fig. 4c,e**) relative to vehicle control across all four brain regions tested, and there was no difference in the number of cells labeled by X-TRAP and TRAP in any of these regions (**Fig. S2**). Moreover, the magnitude of this increase in labeling was similar to the increase in endogenous FOS expression in these brain regions (**Fig. 4d,e**). Thus, X-TRAP and TRAP both recapitulate the broad activation pattern caused by semaglutide injection in the AP, NTS, PBN and CeA.

We next quantified the overlap between the individual cells labeled by X-TRAP, TRAP and endogenous FOS across all four brain regions (**Fig. 4f-i** and **Fig. S2a-d**). This established the following: (1) There was no difference between X-TRAP and TRAP in either their efficiency (%FOS cells labeled by TRAP or X-TRAP) or their specificity (%X-TRAP or TRAP cells labeled by FOS) in any brain region (**Fig. S2a-d**). Thus, the two methods capture the cells that are activated by semaglutide to a comparable extent and with comparable fidelity. (2) A high percentage of X-TRAP cells were also labeled by TRAP (64-81%, **Fig. 4f**) and vice versa (TRAP cells also labeled by X-TRAP: 48-70%, **Fig. 4g**). Thus, there is extensive, but not complete, overlap between the cells labeled by the two approaches. The residual variation likely reflects, in part, the fact that the stimuli were administered on separate days and, therefore, identical brain states were not trapped (see *Discussion*). (3) Almost all of the neurons that were labeled by both X-TRAP and TRAP also expressed endogenous FOS in response to semaglutide (90-95%, **Fig. 4h**). This suggests that cells labeled by both methods are those cells that respond most robustly to the stimulus. (4) 36-56% of all FOS positive cells were doubly labeled by both X-TRAP and TRAP (**Fig. 4i**). This suggests that about half of the cells that express FOS on any given day belong to this robustly activated ensemble.

Overall, these data show that X-TRAP and TRAP label an extensively overlapping population of cells when the same stimulus is administered on different days. Moreover, almost all of these doubly-labeled cells are genuinely activated by the stimulus, as measured by endogenous FOS expression.

### X-TRAP enables visualization of the neurons activated by three stimuli in one animal

One potential application of X-TRAP is to enable visualization of the single-cell overlap between the neurons that respond to three different stimuli (**Fig. 1b**). This could be achieved by using X-TRAP in combination with TRAP to enable labeling for Cre and Flp recombination in two colors, and then staining for endogenous FOS in a third. To test this, we compared the response to semaglutide injection (labeled by TRAP) with the response to lithium chloride (LiCl) injection (labeled by X-TRAP) and consumption of a high-fat diet (HFD; labeled by endogenous FOS).

The rationale for this comparison is that the weight-reducing effects of semaglutide have been proposed to involve two separate mechanisms: the induction of nausea (which is mimicked by LiCl injection) and the induction of “physiologic satiety” (which is mimicked by HFD consumption). However, the extent to which these two mechanisms are mediated by the same or separate circuits is a subject of active controversy (Huang et al., 2024; Yacawych et al., 2025).

We first confirmed that X-TRAP faithfully labels the neurons activated by LiCl injection (**Fig. S3**). We then performed triple labeling in response to semaglutide, LiCl, and HFD (**Fig. 5a**) and analyzed the labeling throughout the brain. We performed this analysis both at the level of region-specific labeling density (**Fig. 5b**) as well as by quantifying the three-way overlap between these markers at the single-cell level (**Fig. 5d-g**). This revealed a number of differences between these stimuli. For example, HFD induced comparatively little labeling in the AP but much more labeling in the BNST, whereas the opposite pattern was observed for semaglutide and LiCl (**Fig. 5b-g**), which is consistent with the fact that the AP is critical for nausea (Zhang et al., 2021) whereas the BNST is broadly involved in ingestion (Canovas et al., 2025). Despite these differences in overall density, we found that there was convergence at the level of individual cells, with most of the AP neurons labeled by HFD also labeled by LiCl or semaglutide (73%, **Fig. 5d**).

**Figure 5:**
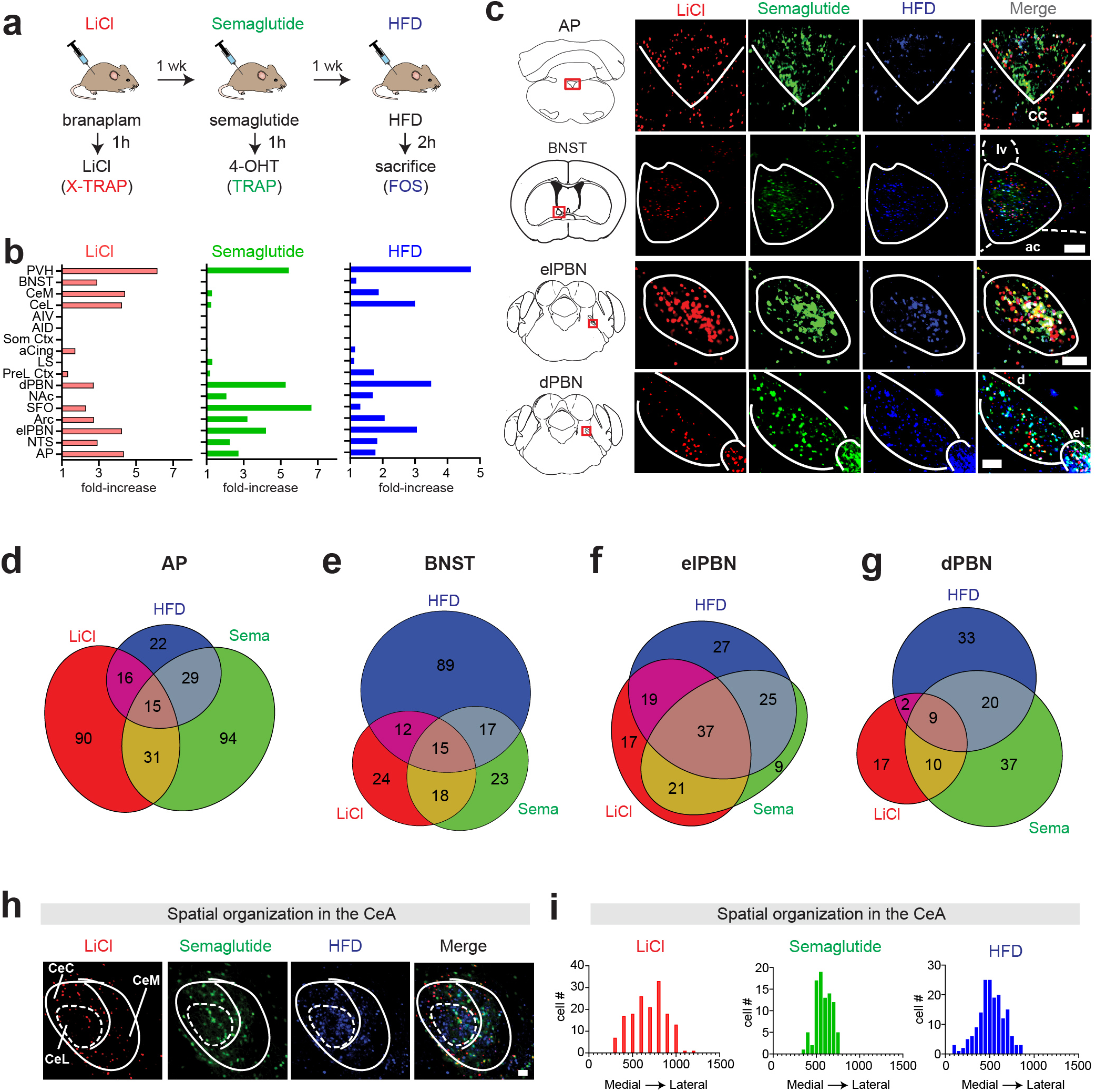
Three-color labelling of neurons activated by LiCI, semaglutide, and HFD consumption. **a.** Protocol for sequential labelling of neurons activated by three drugs. **b.** Number of cells labelled across different brain regions, expressed as the fold-increase relative to vehicle injected controls. Abbreviations: PVH - Paraventricular Hypothalamus, BNST - Bed Nucleus of the Stria Terminalis, CeM - Central Amygdala, medial part, CeC - Central Amygdala, capsular part, CeL - Central Amygdala, lateral part, AIV - Anterior lnsula, ventral part, AID - Anterior lnsula, dorsal part, Som. Ctx - Somatosensory Cortex, aCing - Anterior Cingulate Cortex, LS - Lateral Septum, PreL. Ctx - Prelimbic Cortex, dPBN - Lateral Parabranchial Nucleus, dorsal part, NAc - Nucleus Accumbens, SFO - Subfornical Organ, Arc - Arcuate Hypothalamus, elPBN - Lateral Parabranchial Nucleus, external lateral part, NTS - Nucleus of the Tractus Solitarius, AP - Area Postrema **c.** Examples of labelling in selected brain regions in response to the stimuli indicated. Abbreviations: cc - central canal, ac - anterior commissure, Iv - lateral ventricle. **d-g.** The single-cell overlap between the neurons activated by these stimuli in the brain regions indicated. **h.** Example of labelling within different subdivisions of the CeA. **i.** Quantification of the differential labelling by the three stimuli in the medial to lateral axis. Scale bars are 0.1 mm.

We also observed stimulus-specific differences in overlap within subregions of individual nuclei. Within the PBN, there was extensive overlap at the single-cell level between these three stimuli in external lateral subdivision (elPBN; 90% of semaglutide activated neurons were also labeled by HFD or LiCl; **Fig. 5c,f**). On the other hand, within the adjacent PBN dorsal subdivision (dPBN), there was much less overlap between the three stimuli despite a substantial level of overall labeling (**Fig. 5c,g**). Similarly, within the CeA, HFD caused extensive labeling in the central lateral nucleus (CeL), whereas LiCl largely spared this subnucleus and instead labeled the surrounding capsular (CeC) and medial (CeM) subdivisions (**Fig. 5h**). To quantify these spatial differences, we took advantage of the fact that all three stimuli are labeled on the same brain slice in order to register them on a single coordinate axis and plot the density of labeling across the medial-lateral axis, which revealed distinct distributions (**Fig. 5i**). In a separate experiment, we used this approach to map the overlap between the neurons that respond to three different weight-loss drugs (Hansen et al., 2021). This revealed brain regions where the activated neurons show high overlap (e.g. NTS and AP) and others where they are mostly non-overlapping cells (e.g. somatosensory cortex and SFO) (**Fig. S4**). Overall, these studies show that X-TRAP can be used, in conjunction with TRAP and FOS immunohistochemistry, to reveal the relationship between the responses to three different stimuli at single-cell resolution throughout the mouse brain.

### Generating an AND gate using X-TRAP, TRAP, and an intersectional reporter

We next asked whether we could use X-TRAP and TRAP in combination with an intersectional reporter to deliver a transgene only to cells that respond to two stimuli. We generated by breeding mice that express the X-TRAP and TRAP alleles as well as the intersectional GCaMP7s reporter ai195 (*Fos^XTRAP/TRAP2^ Igs7 ^tetO-GCaMP7s,CAG-tTA2^*). This reporter is designed to express GCaMP7s only after both Cre and Flp recombination. We then treated these animals sequentially with LiCl/branaplam and semaglutide/4-OHT to induce recombination in neurons that respond to these two stimuli (**Fig. 6a**). We also examined two control groups: a drug control (saline injections in place of LiCl/semaglutide) and a genetic control (mice lacking the X-TRAP allele).

**Figure 6.**
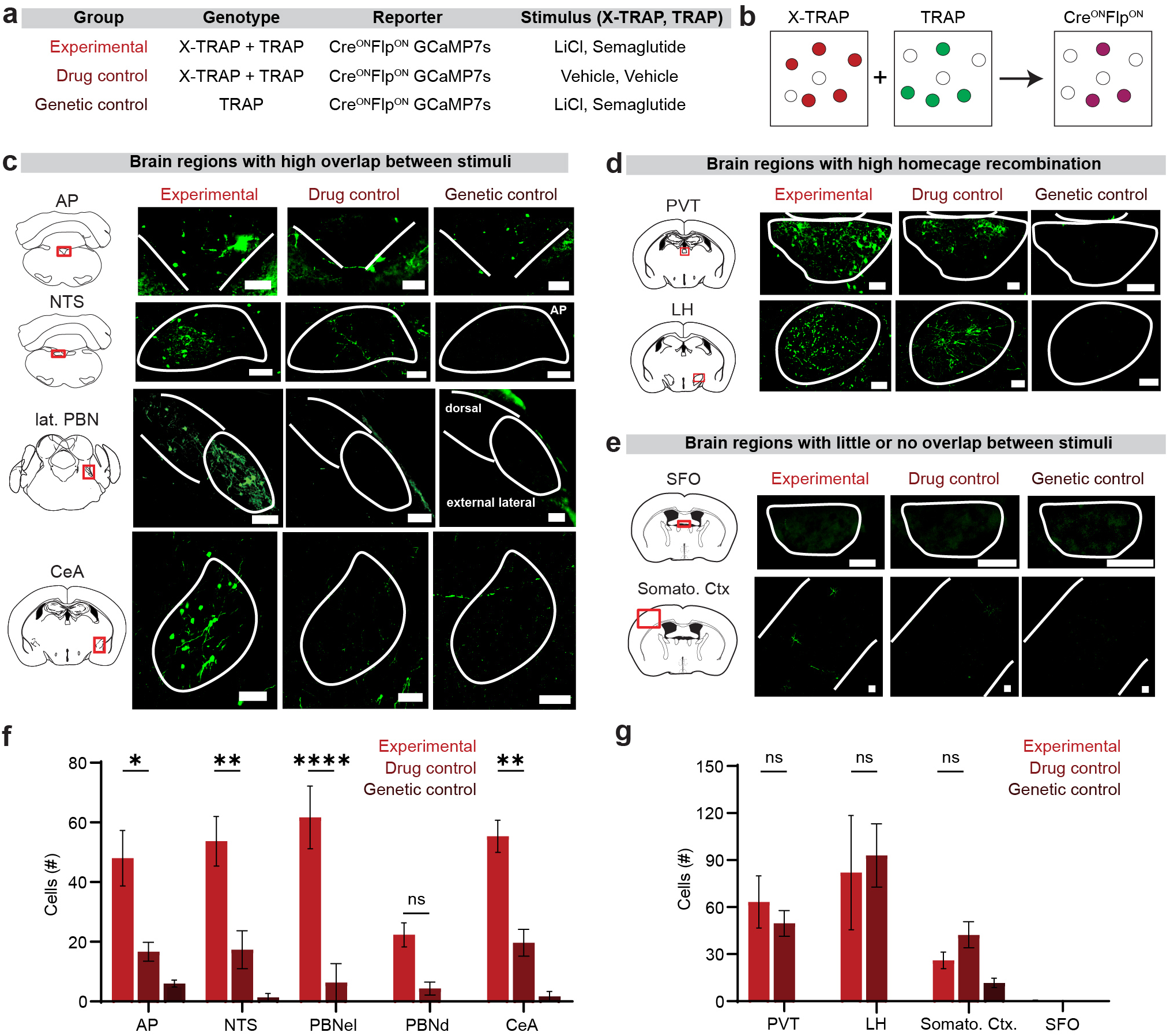
Use of X-TRAP with an intersectional reporter to target GCaMP7s to neurons that respond to two different stimuli. **a.** Description of the experimental cohorts. **b.** Schematic of intersectional Cre-ON Flp-ON labelling. **c.** Expression of GCaMP7s in brain regions that are known to express FOS in response to both semaglutide and LiCI. **d.** Expression of GCaMP7s in brain regions that exhibit high homecage recombination with both TRAP and X-TRAP. **e.** Expression of GCaMP7s in brain regions that do not express FOS in response to both semaglutide and LiCI. **f.** Quantification of the number of cells labelled from each cohort across the brain regions in (c). **g.** Quantification of the number of cells labelled from each cohort across the brain regions in {d,e). Scale bars are 0.1 mm. NS, *P<0.05, **P<0.01, ***P<0.001, ****P<0.0001. Data are mean± sem.

We found that, for the experimental group, there were many cells expressing GCaMP7s in brain regions that respond to both semaglutide and LiCl, such as AP, NTS, PBN and CeA (**Fig. 6c,f**), confirming that this approach can be used to label cells that respond to two different stimuli. In addition, the number of dual labeled cells was larger in the experimental group than in the drug control group, confirming that these are drug-induced responses and not background recombination (e.g. PBNel: 62 ± 10 experimental vs 6.3 ± 6.3 drug control cells, p<0.001). In a few regions such as the AP we observed a small number of GCaMP7s+ cells in the genetic control group (**Fig. 6c,f**), indicating that the ai195 reporter has a small amount of leaky expression.

We also examined brain regions, as the PVT and LH, that we previously found to have (1) high background recombination with both TRAP and X-TRAP (**Fig. 2**), and (2) only moderate FOS expression in response to semaglutide and/or LiCl. As expected, we found that these regions had more cells expressing GCaMP7s in the drug control group, and this was not significantly altered by treatment with semaglutide and LiCl (**Fig. 6d,g**). Finally, we examined brain regions, such as the SFO, that exhibited low background recombination with TRAP and X-TRAP but also did not express FOS in response one of the two drug stimuli (**Fig. 6e,h**). As expected, these brain regions showed minimal GCaMP7s expression. Taken together, these data show that TRAP and X-TRAP can be used in combination with an intersectional reporter to target transgene expression specifically to the cells that respond to two stimuli.

## Discussion

Much recent progress in neuroscience has been driven by the ability to gain genetic access to functional populations of neurons (Luo et al., 2018). While Cre driver mice provide one way to access cell types, the substrates of most brain functions are not defined by a single marker gene, and, for this reason, strategies for activity-dependent genetic labeling have become widely used (DeNardo and Luo, 2017; Franceschini et al., 2020). However, the specificity of these approaches is inherently limited by the fact that they capture all the neurons that are activated during a broad time window surrounding a single stimulus.

To address this problem, efforts have been made to shorten the time window during which activated neurons are labeled. This has been achieved by using calcium, rather than *Fos*, to identify activated neurons and by using light to restrict the temporal window for labeling (Fosque et al., 2015; Lee et al., 2017; Wang et al., 2017). However, these modifications introduce their own problems, such as the need for an invasive fiber optic implant; the restriction of labeling to a small, pre-selected brain region; and the fact that absolute calcium levels do not distinguish tonic from stimulus-induced activity. This has limited the widespread adoption of these approaches.

Here, we have described a conceptually different approach for enhancing the specificity of activity-dependent genetic labeling. This approach is based on using *Fos* to drive the expression of two recombinases regulated by two different small molecules, which enables intersectional labeling of the neurons activated by two stimuli administered on different days. We implemented this by developing a transgenic mouse in which expression of Flp is driven by the *Fos* gene and gated by the small molecule branaplam. We showed this mouse can be used to induce Flp recombination selectively in the neurons that respond to diverse stimuli. We further showed that, when combined with the TRAP allele in the same animal, these two tools enable the simultaneous labeling of neurons that respond to different stimuli with Cre and Flp. This approach should enhance our ability to identify and manipulate the neural correlates of behavior.

### Potential applications of X-TRAP

We envision three ways that X-TRAP and TRAP can be used together to enable new kinds of experiments. First, these tools can be used conjunction with a Cre-On Flp-ON reporter to produce an AND gate in which transgene expression is restricted to the cells that respond to both stimuli. This would be useful when the goal is to access the neurons that encode a common feature of two different stimuli. An example would be two drugs that have different mechanisms of action but elicit the same behavioral response, or two bodily states that are associated with a common physiologic phenotype. We used this intersectional approach to target expression of GCaMP7s to the neurons that are activated by both LiCl, which induces visceral malaise, and semaglutide, which induces malaise but also physiologic satiety (**Fig. 6**).

A second application is to use these tools in conjunction with a Cre-ON Flp-OFF reporter (or Cre-OFF Flp-ON reporter) to produce a NOT gate in which transgene expression is restricted to the cells that respond to one stimulus but not the other. This would be useful when a stimulus has multiple components, one of which can be mimicked by a second stimulus and thereby removed by subtraction. Of note, several lines of Cre-ON Flp-OFF reporter mice have been generated and are available from Jackson laboratory, including lines that express ChR2 (Madisen et al., 2012) as well as the chemogenetic actuators hM3Dq and hM4Di (Zhu et al., 2016) in this way. Thus, it is possible to perform these activity-dependent, subtractive labeling experiments on a brainwide scale.

A third application of these tools is to enable visualization of the single-cell overlap between the neurons that respond to three different stimuli. This can be achieved by performing three-color labeling for Cre and Flp recombination as well as endogenous FOS. We used this strategy to visualize the overlap between the neurons that respond to semaglutide, food ingestion (HFD), and sickness (LiCl) (**Fig. 5**) as well as to visualize the overlap between the neurons that respond three different weight-loss drugs with different mechanisms of action (**Fig. S4**).

### Practical considerations for the use of X-TRAP

There are differences between X-TRAP and TRAP that are important to consider when designing experiments using both tools. The first is that branaplam acts at the level of mRNA, whereas 4-OHT acts at the level of protein. Because *Fos* mRNA is transiently expressed and rapidly degraded, it is important that branaplam reach sufficient concentrations in the brain before the stimulus is administered, so that branaplam can induce alternative splicing of the transgene as it is transcribed. Consistently, we found X-TRAP produced the most efficient labeling when we injected branaplam one hour before the stimulus. On the other hand, the CreER^T2^ protein has a much longer half-life than *Fos* mRNA and likely accumulates over a timescale of at least an hour after stimulus administration. Consistently, we found that TRAP was most efficient when 4-OHT was administered one hour after the stimulus. However, it is important to note that these are rough guidelines, and the optimal injection time will need to be determined empirically in each experiment.

A second consideration relates to the efficiency and specificity of labeling. In general, there was a trend in our data in which TRAP labeled more cells than X-TRAP, which resulted in higher stimulus-induced labeling with TRAP but also higher background. This difference was not apparent in all experiments and probably depends on the strength or duration of the stimulus.

The greater efficiency of TRAP could be due to greater efficiency of Cre versus Flp recombinase (Andreas et al., 2002; Buchholz et al., 1996) or the fact that CreER^T2^ protein likely has a longer half-life than the *Fos*-*Flp* mRNA. Regardless, this observation suggests that it may be beneficial to pair X-TRAP with the stronger stimulus and TRAP with the weaker stimulus when designing dual labeling experiments.

A final consideration regards when to administer the different stimuli. In pilot experiments, we tested injecting mice with branaplam and 4-OHT on the same day, separated by one hour, and we found that this combination resulted in high background recombination, possibly due to some stress associated with injecting these two drugs in close proximity. While it may be possible to reduce this background recombination through habituation, we decided instead to perform only one labeling per day in the experiments described in this study. Despite the fact that branaplam and 4-OHT were injected on different days, usually separated by a week, we found that there was a high degree of overlap between X-TRAP and TRAP recombination when the same stimulus was used (e.g. **Fig. 4**). This confirms that a robust stimulus such as semaglutide reproducibly activates the same neurons on different days. However, it is important to note that there will always be differences in the brain state of the animal when TRAP experiments are performed on different days, and this will result in some variability in the neural ensemble that is labeled.

## Acknowledgements

This work was supported by the National Institutes of Health (R01-DK106399, R01-DK138127, and R01-DK145100 to ZAK). We thank the University of Michigan Pharmacokinetics Core for help with PK experiments. Z.A.K. is an Investigator of the Howard Hughes Medical Institute.

## Methods

Experimental protocols were approved by the University of California, San Francisco IACUC following the National Institutes of Health guidelines for the Care and Use of Laboratory Animals.

### Mouse strains

Experimental animals (>6 weeks old, both sexes) were maintained in temperature- and humidity-controlled facilities with 12 h light-dark cycle and ad libitum access to water and standard chow (PicoLab 5053). We obtained the following mouse strains from Jackson Labs: C57BL/6J (#000664), Fos^CreER/CreER^ (#030323), Ai65F (JAX #032864), Ai195 (#034112), and Ai224 (#037382).

X-TRAP mice were generated by homologous recombination at the endogenous *Fos* locus, aided by targeted CRISPR endonuclease activity. Briefly, to target expression of X-ON:Flp to the *Fos* locus, a targeting construct was synthesized consisting of, from 5’ to 3’: a 2kb homology arm, the X-ON cassette, the FlpO coding sequence, and a 1 kb homology arm. The homology arms were designed such that the X-ON cassette was placed immediately upstream of the *Fos* start codon, and the entire *Fos* amino acid coding sequence (i.e. from the start codon to the stop codon) was deleted and replaced with X-ON:FlpO sequence. The sequence of the X-ON switch was taken from an X-ON Luciferase plasmid (Addgene #174659). To enable insertion of the transgene at the correct genomic site, two sgRNAs were synthesized, one of which directs double strand breaks near the *Fos* start codon (gttgaaacccgagaacatca) and one of which directs double strand breaks near the *Fos* stop codon (gccttctctgactgctcaca). In both cases, the corresponding PAM sequence was mutated in the targeting vector to prevent targeting vector cleavage.

Super-ovulated female FVB/N mice were mated to FVB/N stud males, and fertilized zygotes were collected from oviducts. Cas9 protein (100 ng/uL), sgRNA (250 ng/uL), and targeting vector DNA (100 ng/mL) were mixed and injected into the pronucleus of fertilized zygotes. Zygotes were implanted into oviducts of pseudopregnant CD1 female mice. Screening of the pups by qPCR identified independent founder lines that contained insertion of Flp but not sequences from the targeting vector (i.e. knock-ins). We characterized these founder lines by crossing them to Ai65F reporter mice (JAX #032864), which have Flp-dependent tdTomato expression, and then examining recombination in brain slices +/- branaplam treatment. One line that exhibited highly specific, branaplam-dependent recombination was taken forward for further study.

### Drug preparation for TRAP and X-TRAP

For the X-TRAP experiments, Branaplam hydrochloride (10 mg, MedChemExpress) was dissolved in 1 mL of an 18% solution of 2-hydroxypropyl-beta-cyclodextrin (HP-b-CD, Sigma Aldrich). This 18% solution was prepared by dissolving HP-b-CD (1.8 g) in water (8 mL). The day of experiment, animals were injected with a dose of 25 mg/kg. For TRAP2 experiments, 4-hydroxytamoxifen (4-OHT) (HelloBio HB6040) was dissolved in 200 proof ethanol at a concentration of 20 mg/mL and then shaken at 37°C for ∼60 min (wrapped in aluminum foil) to aid dissolution before being stored at -20°C in 0.25 mL aliquots. On the day of experiment, the dissolved 4-OHT aliquots were shaken at 37°C for 30 min, mixed with corn oil (0.25 mL solution into 0.5 mL corn oil), shaken for an additional 30 min at 37°C, and then the ethanol was evaporated by vacuum centrifugation (SP Genevac, Scientific Products) for 1h. The drug was then injected the same day at a dose of 50 mg/kg.

### Measurements of branaplam pharmacokinetics

Branaplam was administered by IP injection (25 mg/kg) in CD-1 mice. Blood samples were then collected at selected time points (0.083h, 0.167h, 0.25h, 1h, 2h, 4h, 7h, 16h and 24h) using heparinized calibrated pipettes. Samples were centrifuged at 15000 rpm for 10 min, blood plasma was collected from the upper layer, and this plasma was frozen at 80°C for later analysis. The branaplam concentration in these plasma samples was determined by LC-MS/MS by comparing to a standard curve generated by spiking blank plasma with branaplam of known concentrations. Brain concentrations were determined using an analogous approach in which branaplam was administered by IP injection (25 mg/kg) in CD-1 mice, the mice were sacrificed and the brain was rapidly isolated at the indicated times (1h, 2h, 4h, and 16h), and then drug concentration was analyzed by LC-MS/MS. All of these experiments were performed by the University of Michigan Pharmacokinetics Core.

### Drug preparation for experimental stimuli

For the salt-challenge experiments, we prepared a 2 M NaCl solution and administered 250 µL per mouse. For ghrelin experiments, ghrelin was diluted in 0.9% saline (0.2 mg/mL), and mice received a dose of 2 mg/kg. For haloperidol treatment, we prepared a 10 mg/mL stock solution in DMSO, which was diluted on the day of injection (7.5 µL stock into 100 µL saline, yielding ∼0.7 mg/mL), and mice were injected at ∼2 mg/kg. For semaglutide experiments, 3 mg of drug was dissolved in 1 mL DMSO, vortexed, aliquoted (10 µL) and stored at −20 °C. On the day of injection, 3.3 µL of this stock was further diluted into 1 mL of vehicle (0.05% Tween-80 in PBS) to generate a ∼0.01 mg/mL working solution, and mice were injected at ∼0.04 mg/kg. For setmelanotide, a 1 mg/mL stock was prepared in PBS, incubated at 37°C for ∼30 min, aliquoted (500 µL), and stored at −20°C. On injection day, the stock was diluted in 4% bovine serum albumin in PBS to prepare a 0.2 mg/mL solution, and mice were injected at a fixed dose of 2 mg/kg. For rimonabant, a 1 mg/mL stock was prepared in solution of 5% DMSO, 5% Cremophor, 90% PBS and stored at −20°C. On the injection day, mice received a fixed volume of 250 µL. To induce visceral malaise, mice were injected with LiCl at 100 mg/kg in a volume of 250 µL. All vehicle (control) injections were delivered in a fixed volume of 100 µL.

### Experimental timeline

To study branaplam’s effects on feeding and drinking, C57/B6 mice were food or water deprived overnight (18 h; fasted), or provided ad libitum access to food and water (fed), then injected with branaplam or vehicle and 1h later given access to food or water for testing.

For testing the effect of branaplam on background recombination, one group of animals were sacrificed by perfusion without any treatment (branaplam –). The other group was treated with branaplam, and 1h later with 0.9% saline, and then 2 weeks later sacrificed (branaplam +).

For the experiments in Fig. 3, mice were injected with branaplam, and then 1 h later with the stimulus. For the salt challenge, mice were given an injection of 2M salt, the water bottle was removed for three hours, and then mice were returned to their homecage. For the ghrelin injection, mice were given an injection of ghrelin, food was removed for three hours, and then mice were returned to their homecage

For the stimulus-dependent recombination experiments in Fig. 4, 5, and 6, animals were subjected sequential labeling experiments separated by a week. Before the first labeling experiment, mice were food restricted overnight (18 h), then injected with branaplam followed by the stimulus or saline 1h later, and then given re-access to food after six hours. One week later, animals were fasted overnight again, and then injected with stimulus or saline, followed by 4-OHT 1h later, and then given re-access to food 6 h later. For experiments in Fig. 4 and 5, animals were subjected to a third labeling one week later in which they were gain fasted overnight, then refed with HFD or injected with stimulus/saline, and then perfused 2h later to enable staining for endogenous Fos.

### Histology

Animals were anaesthetized under isoflurane and then transcardially perfused with 1XPBS followed by 10% formalin. Brains were dissected, post-fixed in 10% formalin overnight at 4C and switched to 15% sucrose the next day and to 30% sucrose the following day for cryo-protection and embedded in OCT before sectioning. Sections (40 µm) were prepared using a cryostat and collected in 1xPBS and stored at -4°C wrapped in aluminum foil for protection from light. For immunostaining, sections were washed 3 × 10 min with 0.1% PBST (0.1% Triton X-100 in PBS), blocked (2% NGS in 0.1% PBST) for 1h at room temperature and then incubated with primary antibodies for ∼48 h at 4°C. After two days, sections were washed 3 × 10 min with 0.1% PBST, incubated with secondary antibodies for 1 h at room temperature, washed again 3 × 10 min with 1 x PBS and mounted with DAPI Fluoromount-G (Southern Biotech).

For Fos staining, rabbit anti-Fos (Abcam, ab190289, 1:1,000 in 2% Normal Goat Serum - NGS) was used as the primary antibody and goat-anti-rabbit 647 (Invitrogen, ab359307, 1:1,000) as the secondary antibody. For the Fos+GFP staining, we used chicken anti-GFP (AvesLab, ab917979, 1:5000) and rabbit-anti-Fos (1:1,000 dilution) in a 2% NGS solution. For secondary staining, we used chicken anti-GFP 488 (Invitrogen, ab2566343, 1:1,000) and goat anti-rabbit 647 (Invitrogen, ab359307, 1:1,000) in 2% NGS. For the GCaMP7 staining, anti-GFP (AvesLab, ab917979 1:5,000 in 2% NGS) were used as a primary and goat anti-chicken 555 (Invitrogen, ab1964371, 1:1,000 in 2%NGS) as a secondary.

### Imaging and QUINT analysis

Sections were imaged at 10x using a Nikon Eclipse Ti2 widefield microscope with consistent exposure and gain. Images were exported as TIFF files and were manually quantified in Fiji. For unbiased brain-wide labeling shown in F2d and F3c, whole-slide imaging of brain tissue was performed using NIS Elements with a custom JOBS. Brain wide Tomato+ expression analysis to identify FLP recombination in response to diverse stimuli was done using QUINT (https://quintworkflow.readthedocs.io/en/latest/QUINTintro.html)^#^ (**Fig. 3**). Section images were curated and preprocessed using *Nutil* tools (*TiffCreator, Transform, Resize*) to ensure compatibility with the QUINT pipeline. Images were registered to the Allen Mouse Brain Atlas 2017 in a two-step procedure. First, each section was manually aligned to the reference atlas using *QuickNII.* This was followed by in-plane refinement of the alignment using *VisuAlign,* which applies non-linear registration methods to improve anatomical correspondence. Following registration, images were imported into *Ilastik* for image segmentation. Segmentation was performed in two stages: (1) *Pixel Classification* to distinguish immunoreactive signal from background, and (2) *Object Classification* to separate real cells from artifacts. A subset of sections of different experiments (2M salt, ghrelin, haloperidol, saline) was used to train both classifiers, which were subsequently applied to the entire dataset in batch mode, greatly reducing processing time relative to manual segmentation. Registered and segmented images were then combined to generate atlas-mapped datasets, which were imported into Nutil Quantifier. This enabled automated, brain-wide quantification of immunoreactivity across anatomical regions defined by the Allen atlas.

### Statistical Analysis

Graphing and analyses were performed in GraphPad Prism 10 (Graph Pad Software). P values for comparisons across groups were calculated using the one-way ANOVA test and corrected for multiple comparisons using Tuckey’s multiple comparisons test. All values are reported as the mean ± s.e.m. (error bars). For Figs. 1F–I, paired t-tests were used to assess within-subject differences in feeding or drinking behavior following branaplam versus saline treatment in the same animals. In figures, asterisks denote statistical significance: *\*P* < 0.05, *\*\*P* < 0.01, *\*\*\*P* < 0.001, *\*\*\*\*P* < 0.0001.

---

**Supplementary Figure 1.**
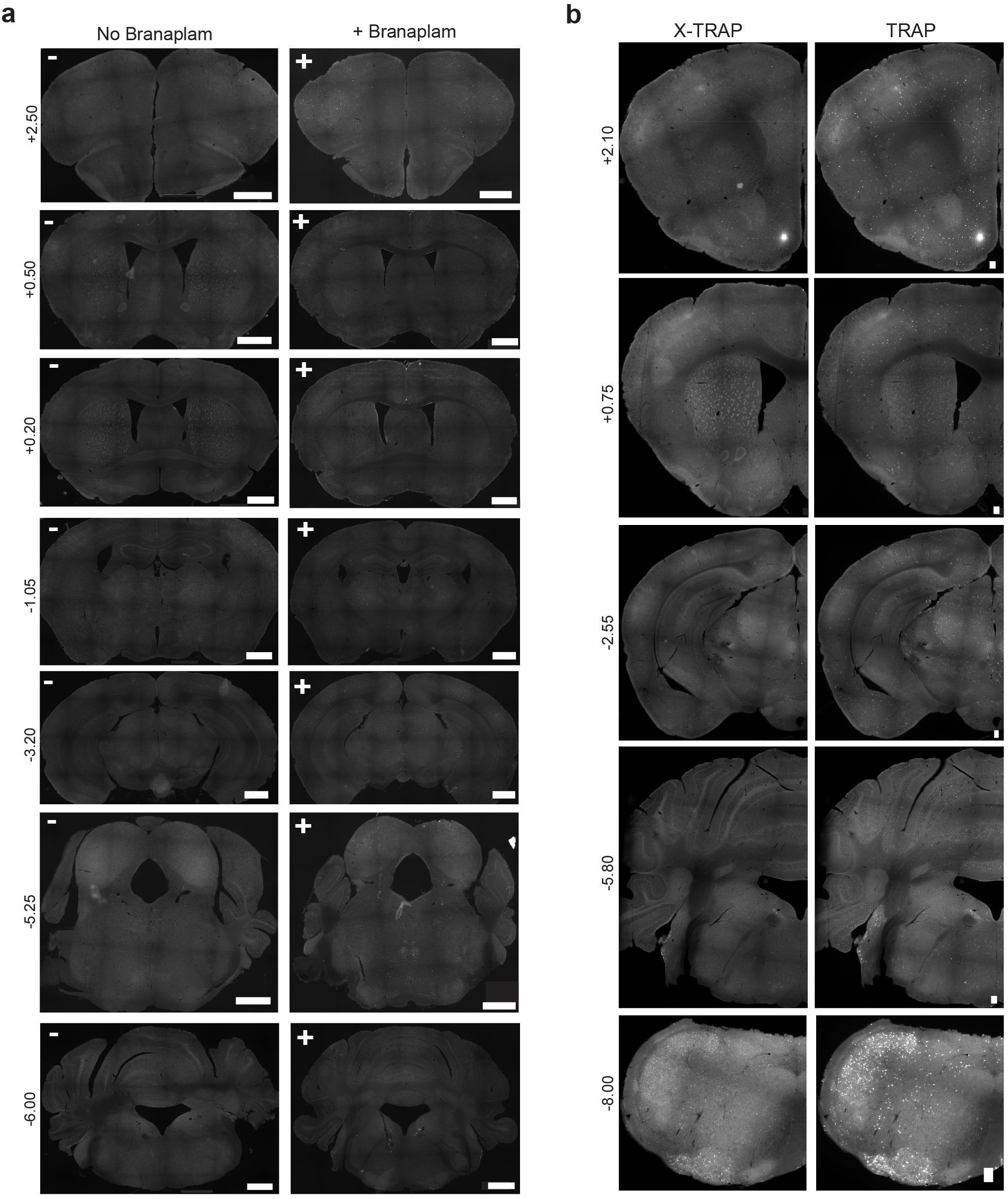
Homecage recombination by X-TRAP and TRAP. **a.** Homecage recombination without (left) and with (right) branaplam treatment. Shown is Tomato fluorescence in an X-TRAP mouse crossed to a Flp-Tomato reporter. Note there are no cells without branaplam, and only sparse recombination after branaplam.The numbers at left are the distance from bregma of the slice. Scale bars are 1 mm. **b.** Homecage recombination with X-TRAP (left) versus TRAP (right) in the same mouse. Note that there is generally less recombination with X-TRAP. Scale bars are 0.1 mm.

**Supplementary Figure 2.**
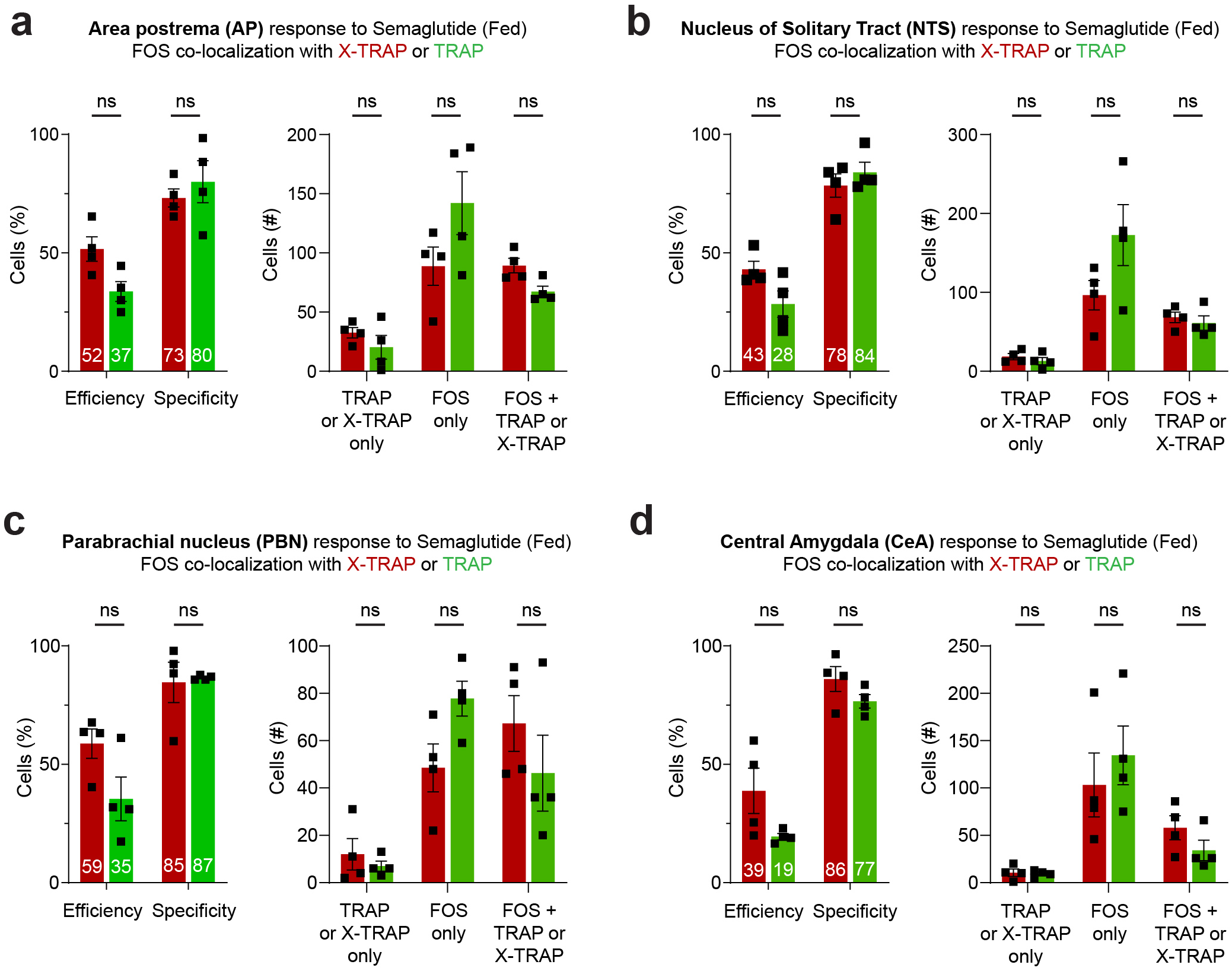
X-TRAP and TRAP label semaglutide-activated neurons with similar specificity and efficiency in fed mice. a-d. Left. Shown in each panel is the efficiency (i.e.% FOS+ cells that are also labelled by X-TRAP (red) or TRAP (green)) and specificity (i.e.% X-TRAP (red) or TRAP (green) cells that are also labelled by FOS). **Right.** Shown in each panel is the number of cells with each labelling. Note that, in this experiment, the X-TRAP and TRAP labelling were performed in different mice. Thus, for example, the red bar in the "FOS only" column is the number of FOS+ cells that were not labelled by X-TRAP.

**Supplementary Figure 3.**
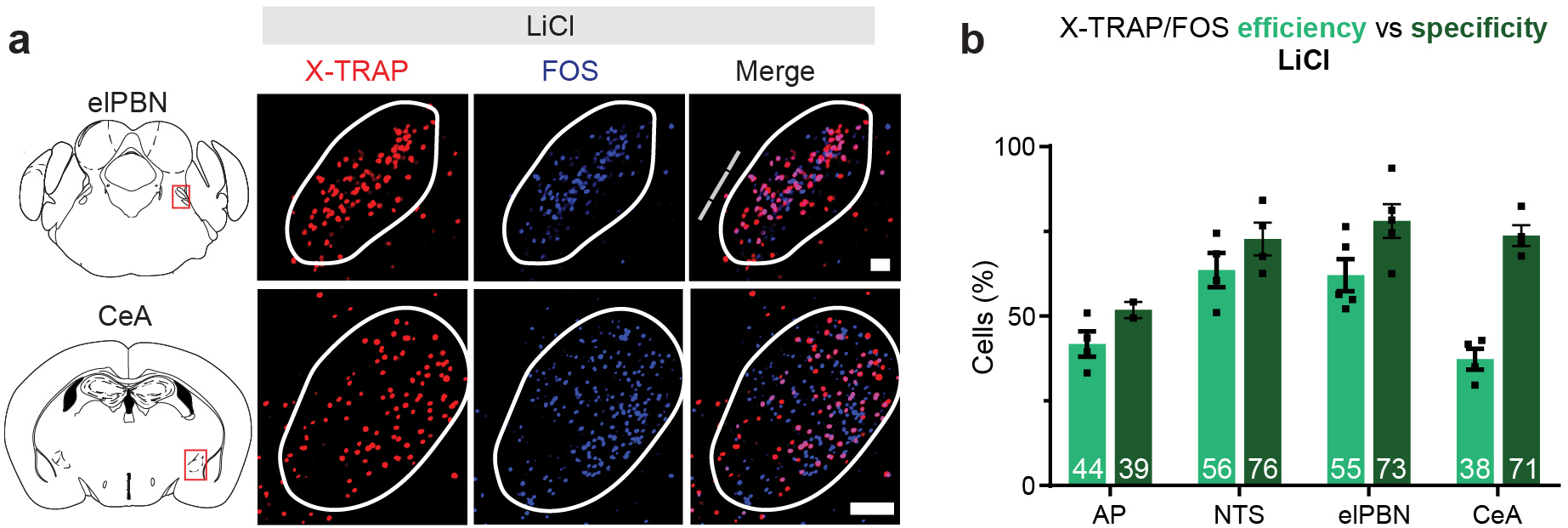
Labeling of LiCI activated neurons by X-TRAP. **a.** Co-localization between X-TRAP recombination and endogenous FOS induced by LiCI treatment. **b.** Efficiency (percentage of FOS+ cells that are Tomato+) and specificity (percentage of Tomato+ cells that are FOS+) for LiCl-induced labelling. Scale bars are 0.1 mm. Data are mean± sem.

**Supplementary Figure 4:**
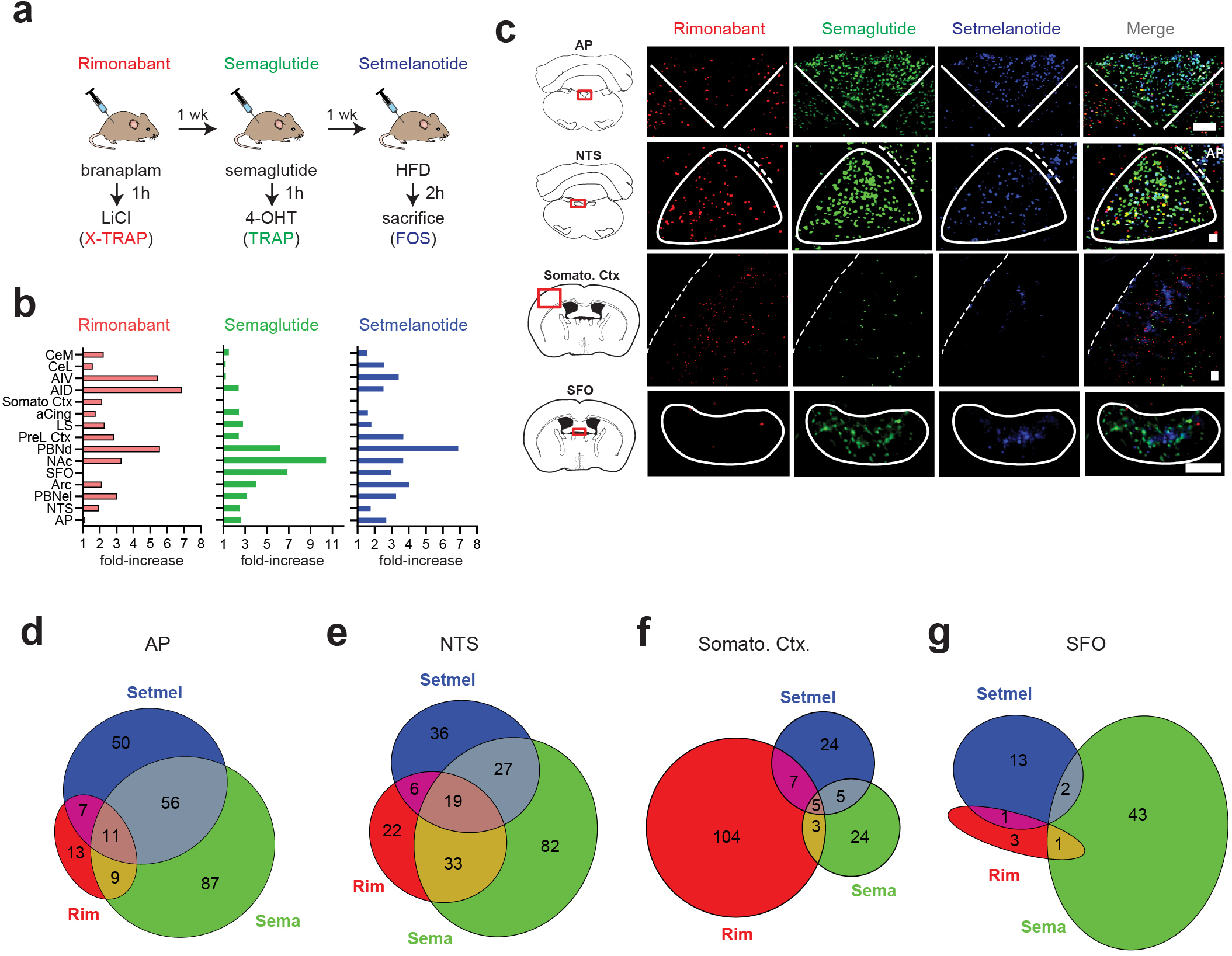
Mapping brainwide responses to weight-reducing drugs. **a.** Protocol for sequential labelling of neurons activated by three drugs. **b.** Number of cells labelled across different brain regions, expressed as the fold-increase relative to vehicle injected controls. Abbreviations: PVH - Paraventricular Hypothalamus, BNST - Bed Nucleus of the Stria Terminalis, CeM - Central Amygdala, medial part, CeL - Central Amygdala, lateral part, AIV - Anterior lnsula, ventral part, AID - Anterior lnsula, dorsal part, Som. Ctx - Somatosensory Cortex, aCing - Anterior Cingulate Cortex, LS - Lateral Septum, PreL. Ctx - Prelimbic Cortex, dPBN - Lateral Parabranchial Nucleus, dorsal part, NAc - Nucleus Accumbens, SFO - Subfornical Organ, Arc - Arcuate Hypothalamus, elPBN - Lateral Parabranchial Nucleus, external lateral part, NTS - Nucleus of the Tractus Solitarius, AP - Area Postrema. **c.** Examples of labelling in selected brain regions in response to drug injection. **d-g.** The single-cell overlap between the neurons activated by these drugs in the brain regions indicated. Scale bars are 0.1 mm.

